# Firing rate distributions in spiking networks with heterogeneous connectivity

**DOI:** 10.1101/542456

**Authors:** Marina Vegué, Alex Roxin

## Abstract

Meanfield theory for networks of spiking neurons based on the so-called diffusion approximation has been used to calculate certain measures of neuronal activity which can be compared with experimental data. This includes the distribution of firing rates across the network. However, the theory in its current form applies only to networks in which there is relatively little heterogeneity in the number of incoming and outgoing connections per neuron. Here we extend this theory to include networks with arbitrary degree distributions. Furthermore, the theory takes into account correlations in the in-degree and out-degree of neurons, which would arise e.g. in the case of networks with hub-like neurons. Finally, we show that networks with broad and postively correlated degrees can generate a large-amplitude sustained response to transient stimuli which does not occur in more homogeneous networks.

## I. INTRODUCTION

One of the aims of neural science is to understand how the observed patterns of neuronal activity originate from the properties of single neurons and the interactions between them. Many experimental studies have revealed that cortical neurons tend to spike at low rates and in a highly irregular manner [1–3], with coefficients of variation (CVs) of inter-spike-intervals (ISIs) in the range 0.5-1 [1]. The current hypothesis to explain the emergence of such irregularity is that, under physiological conditions, neurons receive a large number of both excitatory and inhibitory inputs which cancel in the mean, in such a way that the membrane voltage resembles a random walk between the resting and the threshold potentials. Although the number of total inputs received can be large, the spiking process, that results from the voltage crossing the threshold potential, becomes highly irregular [4]. Such a balanced state has been successfully reproduced in models of neuronal networks without the need for fine-tuning of parameters [5, 6].

In the majority of such studies, network structure is either homogeneous (that is, every neuron receives the same number of connections from the network) or simply random (i.e., connections among neurons are created independently with a fixed probability, as in the so-called Erdös-Rényi (ER) model). However, the structure of cortical microcircuits deviates from these scenarios, as has been directly evidenced in patch clamp experiments performed on brain slices [7–12]. Such observations are based on different measures (such as two- and three-neuron motifs counts, the total number of connections in small neuronal subsets and the connection probability as a function of the number of common neighbors, among others), and are collectively referred to as the “nonrandomness” of the cortical microcircuitry. It remains an open question to what extent the dynamics exhibited by networks of spiking neurons is affected by more realistic topologies.

One of the characteristic properties of homogeneous and simple random networks is that they exhibit little structural heterogeneity: the distribution of in- and out-degrees in the network is tightly peaked around the mean value. Although the degree distribution is not a local property and is therefore very difficult to estimate in a real network without the knowledge of the complete structure [13], many alternative –and more heterogeneous-degree distributions are in principle possible in cortical circuits. Individual pyramidal neurons in cortex exhibit distinct amounts of coupling to the overall activity in the surrounding network [14].

This heterogeneity is, moreover, related to the likelihood of their receiving connections from neighboring pyramidal cells: neurons with high coupling tend to receive more connections from the local network than those with a low level of coupling. Thus, diversity in population coupling might be a functional consequence of structural degree heterogeneity in the circuits of cerebral cortex.

Recent work [15] has revealed that anatomical reconstructions of L4 rat barrel cortex [16] exhibit an important heterogeneity in input connectivity. In previous work [13] we showed that some of the nonrandom features of real cortical circuits are compatible with networks that are defined through broad in/out-degree distributions which are also positively correlated, although such configurations are unlikely in light of other local measures. In any case, the analysis of the structure in small groups of excitatory neurons revealed that in- and out-degrees are positively correlated, a feature that simple random models cannot reproduce [13]. Therefore, it is of particular interest to study the role that broad degree distributions and degree correlations might play in neuronal dynamics. The effect of broadening in- and out-degree distributions in networks of spiking neurons has been studied in [17], in which it was shown that the variability of in-degrees has an important impact on the dynamical state of the network, including global oscillations, whereas the out-degree variance shapes pairwise correlations in the synaptic currents. Broad excitatory distributions of in-degrees can also break down the balanced assumption in model networks unless proper compensatory mechanisms are introduced, such as degree correlations [15, 18], adaptation currents [15] or tuning of the inhibitory weights through synaptic plasticity mechanisms [15].

Apart from marginal degree distributions, degree correlations might play an important role in neuronal activity. Correlations between in-degrees and out-degrees of individual neurons have been shown to substantially affect the stability of binary neural networks [19]. Aljadeff et al. [20] computed the eigenvalues of connectivity matrices in which the excitatory subnetwork exhibits general degree distributions, and showed that positive in/out-degree correlations have an important impact in the spectrum. Nykamp et al. [21] analytically studied the role of such correlations in firing rate models and showed that a positive correlation between excitatory in- and out-degrees has a similar effect to increasing the excitatory weights in the network.

A precise, macroscopic description of networks of leaky integrate-and-fire (LIF) neurons in the balanced state is provided by mean-field theory introduced by Amit and Brunel [22–24]. This theoretical framework can explain some observed features of neuronal dynamics –such as the skewed rate distributions found *in vivo* [2, 3]- in model networks with purely random connectivity [25, 26]. Schmeltzer et al. [27] used a mean-field extension to show that assortative in-degree correlations improve the sensitivity to weak stimuli in model networks (in network theory, the term *assortativity* is used to denote the property by which connected nodes tend to have similar degrees [28]). However, a generalization of such a mean-field formalism to networks of spiking neurons with connectivity structures defined by arbitrary degree distributions is, to our knowledge, still lacking.

In this paper we address the problem of extending such mean-field techniques to networks of excitatory (E) and inhibitory (I) neurons with a highly heterogeneous structure. In particular, we consider the case in which in- and out-degrees follow a prescribed joint distribution, which could include correlations between individual degrees. The extended system of self-consistent equations provides a means to compute the distribution of firing rates in the stationary state. Our results show good agreement between theory and simulations. We use the derived equations to demonstrate that a positive correlation between in- and out-degrees can have important consequences for dynamics, mainly because it biases the firing rate distribution in the set of available pre-synaptic neurons. This effect has been already pointed out by Nykamp et al. [21] in the case of firing rate models.

The presence of broad E-to-E degree distributions can destabilize dynamics and destabilize the stationary state, as long as inhibitory connections are created with a fixed probability. This is due to the fact that in such networks there are neurons which receive a large amount of excitation which is not balanced by inhibition. Heterogeneity in the total amount of excitation received has been observed in the rodent cortex, where compensatory mechanisms at the level of inhibitory synapses are able to maintain a proper balance [29]. We have mimicked such a compensation by allowing inhibitory connections to appear with a probability that is modulated by the total excitatory in-degree. Under such circumstances, the network can return to an asynchronous stationary state, very similar to those exhibited by purely random networks. Interestingly, in networks whose degrees are, in addition, positively correlated, transient external inputs can destabilize the stationary state for a period larger than the duration of the stimulus. This finding suggests a possible role of the degree correlation in enhancing the ability of neuronal networks to respond to transient stimulation.

## II. STANDARD MEAN-FIELD THEORY FOR SPIKING NEURONS

### A. The model

We consider a network of *N* spiking neurons. An arbitrary neuron *i* in the network has a membrane voltage *V*_*i*_ which evolves according to

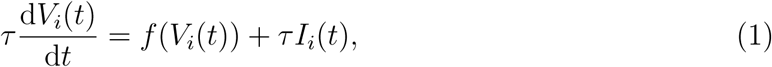

where *τ* is a time constant and *I*_*i*_(*t*) is the synaptic input. Every time *V*_*i*_ reaches a threshold *V*_*θ*_, the neuron generates an action potential and the voltage is immediately reset to *V*_*r*_, where it remains for a resting period *τ*_*r*_. Eq. (1) describes the class of so-called integrate-and-fire neuron models, which are distinguished by the particular form of the function *f* (*V)*. In this manuscript we will analyze networks of leaky integrate-and-fire neurons (LIF) for which *f* (*V)* = –*V*. Nonetheless, the theoretical framework is valid for other choices of neuron model.

The input is generated from the spikes of the pre-synaptic neurons to our neuron. We impose that every action potential emitted by the *j*-th pre-synaptic neuron induces an instantaneous jump in the voltage *V*_*i*_, of magnitude *J*_*ij*_. This is equivalent to saying that the instantaneous variation of *V*_*i*_ at time *t* induced by a pre-synaptic spike emitted by neuron *j* at time *t′* is *J*_*ij*_ *δ*(*t – t′ – d*_*j*_), where *δ* denotes the Dirac delta function and *d*_*j*_ is a synaptic delay associated with neuron *j*. Thus, the external input can be expressed as

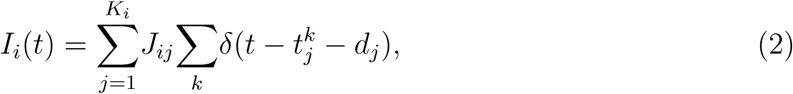

where *K*_*i*_ is the number of incoming connections or in-degree of the neuron under study and 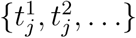 are the spike times of the pre-synaptic neuron *j*.

We are interested in studying macroscopic properties of this system when we impose a certain topology in the network. The first assumption to make is that individual neurons fire as Poisson processes, so that Eq. (2) is non-deterministic and Eq. (1) becomes a *stochastic* differential equation.

### B. Mean-field equations

Classical mean-field analysis provides tools for predicting some statistical properties of the stationary state (such as the firing rates) in networks of this type when the underlying structure is homogeneous or simply random [24–26]. The results are different depending on the topology of the network. In a homogeneous scenario (i.e., when all the neurons are equivalent and have the same in-degree), the stationary state is characterized by a single stationary firing rate. In the random or Erdös-Rényi (ER) case, the variability in terms of in-degrees translates into a variability of stationary firing rates and the stationary state is described by a *distribution* of firing rates.

The necessary conditions for the analysis to be correct are:

(i) neurons fire as independent Poisson processes;
(ii) the sizes of the voltage jumps {*J*_*ij*_}_*i,j*_ are small compared with the threshold *V*_*θ*_ so that the voltage can be approximated by a continuous variable.

Condition (i) is approximately fulfilled when the synaptic input is sub-threshold (which induces irregular spiking) and there is a small overlap in the total input received by any pair of neurons (which ensures independence between inputs to different neurons). A small overlap occurs when the connectivity is random and sparse, that is, when the in-degrees are small compared with the system’s size *N*. Condition (ii) depends on the parameter’s choice and can therefore be assumed in general. From now on, we suppose that conditions (i) and (ii) are fulfilled.

Under these assumptions, the input to a given neuron can be written

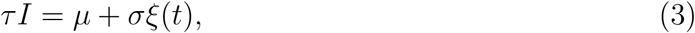

where *µ* and *σ*^2^ are the mean and the variance of the total input received during a time window of length *τ* and *ξ*(*t*) is a Gaussian white-noise process of unit variance. The stochastic evolution of the membrane voltage *V* of a single neuron can be described by means of a Fokker-Planck equation. The steady state solution of the Fokker-Planck equation gives the stationary probability density of the membrane potential. From this one can calculate the stationary probability flux at threshold, which is just the firing rate of the neuron. It is a function of the form *v* = *ϕ*(*µ, σ*). For the case of LIF neurons, the function can be found analytically, and is

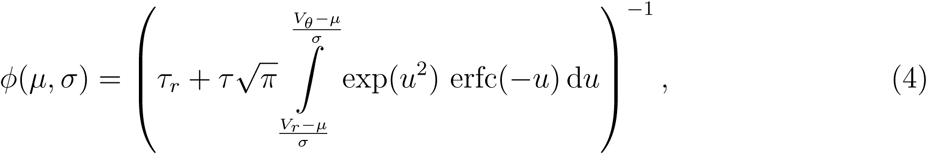

(see ref. [24] for details).

#### Fully homogeneous connectivity

When the network is composed of excitatory (E) and inhibitory (I) neurons with the same connectivity and dynamical properties and the E (I) weights are *J*_*E*_ (*-J*_*I*_), all the neurons are statistically equivalent and their firing rate in the stationary state *v* is the same. Specifically, the input current to a neuron *i* can be written

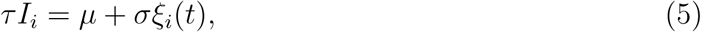

and so the mean and variance are identical for all neurons, while the fluctuations are independent. By *same connectivity properties* we mean that all the neurons receive the same number *K*_*E*_ of E connections and the same number *K*_*I*_ of I connections but the precise realization of this connectivity is totally random and therefore uncorrelated from neuron to neuron; this is what leads to the temporal fluctuations being independent. If each neuron also receives external inputs from an independent set of *K*_ext_ neurons, through synaptic weights *J*_ext_, which fire at a constant rate *v*_ext_, the quantities *µ* and *σ*^2^ depend on *v* through

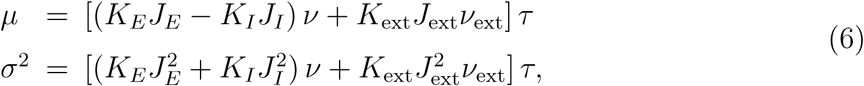

so (4) and (6) define a self-consistent equation for the stationary firing rate *v*.

#### Erdös-Rényi (ER) connectivity

When connections are generated independently with a fixed probability *p* (simple random scenario), neurons are heterogeneous in terms of their in-degrees, and this induces a heterogeneity in the stationary firing rates. In this case, the input to a neuron *i* can be written

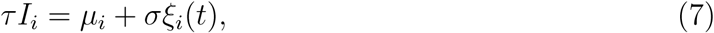

where *µ*_*i*_ (which varies from neuron to neuron) takes the form

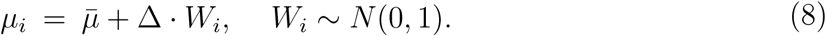

*W*_*i*_ captures both the variability coming from the differences in in-degrees and the variability due to the fact that pre-synaptic neurons fire at different rates. In the limit of a large network, both levels of variability can be put together under this common continuous random variable *W*_*i*_ (see [25, 26] for details).

Briefly, and for identical E and I populations, 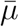 and *σ* are given by Eq. (6) once the degrees *K*_*E*_*, K*_*I*_ have been replaced by their network averages, ⟨ *K* ⟩ _*E*_ = *pN*_*E*_, ⟨ *K* ⟩ _*I*_ = *pN*_*I*_, and *v* has been replaced by the average firing rate in the network,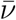 :

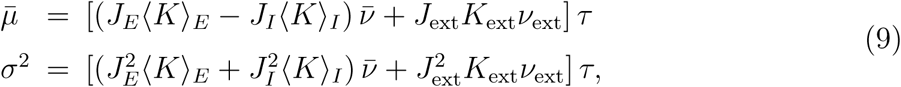

whereas

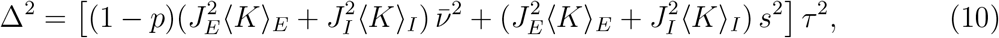

where *s*^2^ is the variance of the firing rate distribution in the network.

Therefore, the stationary firing rates depend on 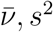 and are parametrized by a standard Gaussian variable *W* through 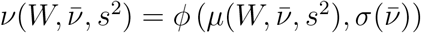. The mean and variance of the firing rate distribution are found self-consistently through the following integrals

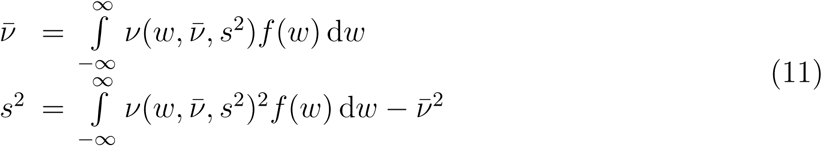

where *f* is the probability density function of a standard Gaussian random variable. Eq. (11) constitutes a system of self-consistent equations for 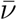 and *s*^2^. If, in addition, E and I neurons are different in terms of other properties, the equations are analogous but depend on the mean and variances of E and I firing rate distributions. In this case the system to be solved has four unknowns and four equations.

In any case, once the corresponding means and variances of the firing rate distribution are calculated, one can reconstruct the entire firing rate distribution *p*(*v*). This is done by passing the input distribution through the steady-state nonlinearity. Formally, one can write

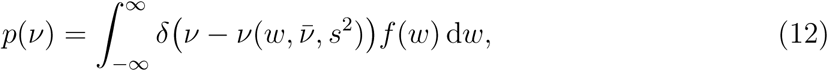

where *δ*(*x*) is the Dirac-delta function (notice that here we are making some abuse of notation: *v* is a number and *v*() denotes the rate function given by 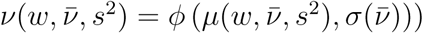. By denoting 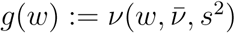, defining the variable *z* = *g*(*w*) and integrating with respect to *z*, Eq. (12) can be written

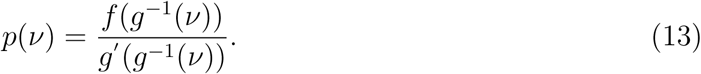

This formula assumes that the steady-state fI curve 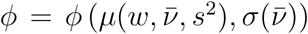 is invertible with respect to *w*.

## III. MEAN-FIELD DESCRIPTION OF NETWORKS WITH ARBITRARY DEGREE DISTRIBUTION

The aim now is to extend the formalism sketched in the previous section to more general classes of networks. We focus here on networks whose connectivity (at least within the EE subnetwork) is generated to preserve a given in/out-degree distribution, as in the Degree model described in [13]. The distributions used will be broader than those provided by ER networks, that is, we consider distributions for which the mean degree ⟨ *K* ⟩ is large when *N* is large but whose standard deviation-to-mean ratio 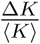 stays constant and it is not necessarily small. We also introduce correlations between individual in- and out-degrees and show how these correlations shape the distribution of firing rates in the stationary state. All the results apply also to the previously studied ER cases.

### A. Networks with arbitrary and independent in- and out-degree distributions

We start by considering the case of networks with arbitrary degree distribution, assuming that the out-degrees are independent of in-degrees. To make the presentation clearer, we assume first that the network is composed of neurons of a single type with homogeneous synaptic weights *J*.

Let us consider a single neuron. Its firing rate in the stationary state is *v* = *ϕ*(*µ, σ*), where *ϕ* is defined by Eq. (4) and *µ, σ* are the mean and the standard deviation of the total input received by the neuron within a time window of length *τ*. We start by computing *µ* and *σ*^2^.

Let us suppose that our neuron receives inputs from *K* pre-synaptic neurons. Since the pre-synaptic neurons fire as Poisson processes, if we knew their rates, *v*_1_*, …, v*_*K*_, these quantities would be

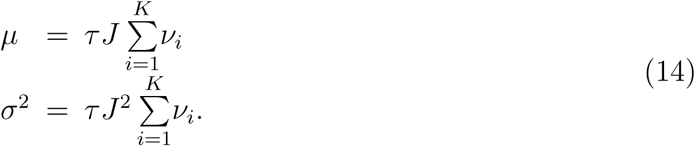

As different neurons have different degrees, we expect to have a *distribution* of firing rates in the stationary state in the entire network. Therefore, the term 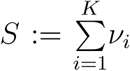 varies across neurons due to differences in their connectivity. If the neuron under study is randomly chosen, *S* will be a random variable that is defined as a sum of independent variables that come from a common firing rate distribution. If *K* is large enough, the Central Limit Theorem ensures that *S* will approximately follow a Gaussian distribution with mean 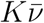 and variance *Ks*^2^, where 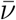 and *s*^2^ are the mean and variance of the stationary firing rate distribution:

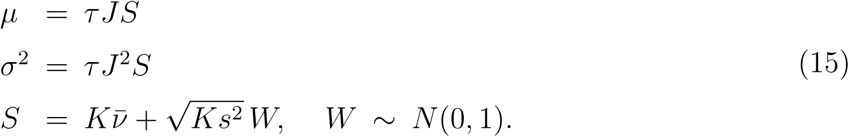

Recall that one of our assumptions is that the network is sparse, meaning that *K ≪ N*. But we also suppose that the degrees are large quantities when *N* is large. In this case, we can approximate *σ* by its leading term 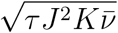.

Regarding now *K* as a variable which varies from neuron to neuron, we can express *µ* and *σ* as a function of the *quenched* randomness given by the pair (*K, W)* as

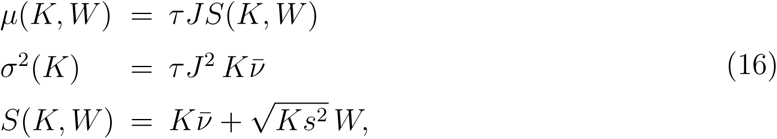

and the stationary firing rate of a neuron with *K* = *k, W* = *w* given the statistics 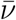 and *s*^2^ of the firing rate distribution is

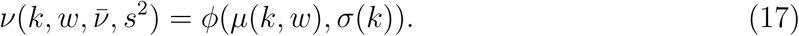

In the large *N* limit we can treat the degrees as continuous random variables. In this case, as in (11), the system of equations can be closed through the definition of 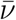 and *s*^2^:

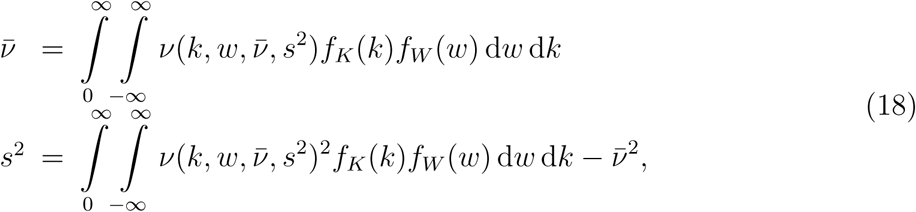

where *f*_*K*_ and *f*_*W*_ are the probability density functions of the *K* and *W* variables. Since *W* is a standard Gaussian, 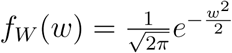. Notice that the integration limit of the variable *K* is set to (0*, ∞*) even though the degree is always bounded by *N* – 1; we assume that the support of *f*_*K*_ is contained in [0, *N* – 1]. To find the distribution of firing rates in this system, we (numerically) solve Eq. (18) for the unknowns 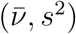. The stationary firing rate of a neuron with *K* = *k* and *W* = *w* is then just obtained by evaluating Eqs. (16), (17). In order to reconstruct the full firing rate distribution we can formally write

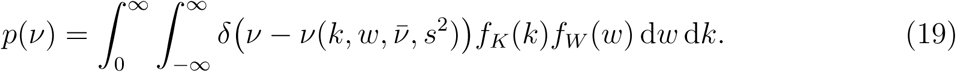

This integral can be further simplified by integrating over the Gaussian variability as in Eq. (12). The resulting formula still requires integration over the in-degree variability.

This formulation can be easily extended to the case of a network composed of E and I neurons where the connectivity rules are different depending on the neuronal type and there is an external source of inputs. Imagine that the external inputs come from a population of neurons which fire as Poisson processes at constant rate *v*_ext_, in such a way that each neuron receives information from a fixed number *K*_ext_ of these external sources, which are totally uncorrelated. We assume that all the E (I) weights are the same and they take the value *J*_*E*_ (*-J*_*I*_). If *K*_*αβ*_ is the random variable which gives the number of inputs from population *β* to a neuron within population *α*, expression (16) reads

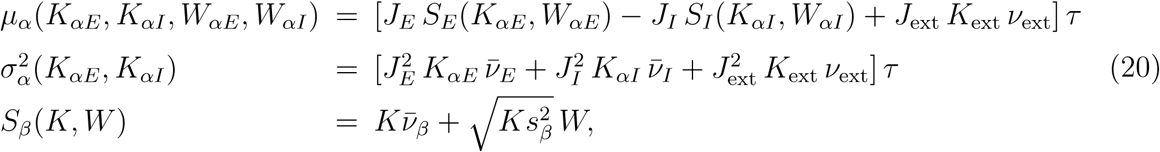

where *μ*_*α*_ and 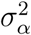 denote the *μ* and *σ*^2^ variables associated with population *α ∈* {*E, I*}, and 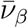 and 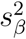 refer to the mean and variance of the distribution of stationary firing rates within population *β*.

Expressions (20) specify the magnitude of *μ*_*α*_ and 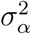 as a function of the random variables *K*_*αE*_, *K*_*αI*_, *W*_*αE*_, *W*_*αI*_. *K*_*αE*_ and *K*_*αI*_ represent the heterogeneity in terms of in-degrees, whereas *W*_*αE*_ and *W*_*αI*_ are normally distributed and reflect the variability in the firing rates of the input neurons. In general situations *K*_*αE*_, *K*_*αI*_, *W*_*αE*_, *W*_*αI*_ will be pairwise independent, except, maybe, for the two “structural” variables *K*_*αE*_, *K*_*αI*_, which could be correlated depending on the connectivity imposed (as, for example, in a network where E and I degrees compensate each other to achieve balance). The expressions for *μ*_*α*_ and 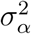 also depend on the mean and variance of the distributions of stationary firing rates in the E and I populations. Therefore, the firing rate of a neuron is a function of *K*_*αE*_, *K*_*αI*_, *W*_*αE*_, *W*_*αI*_, 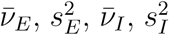 and can be recovered when the mean and standard deviation of the firing rate distributions are known. These quantities can be computed by solving the corresponding extended version of Eq. (18).

#### Comparison with Erdös-Rényi network

In an Erdös-Rényi network, connections are made with a fixed probability *p*, leading to a Binomial distribution of degrees with mean ⟨*K*⟩_*α*_ = *pN*_*α*_ and variance 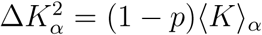 for *α ∈* {*E, I*}. In this case, Eq. (20) can be written

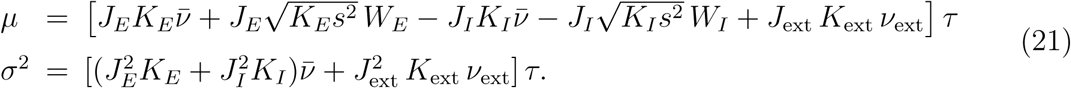

As long as *K*_*α*_ ≫ 1 the degree distribution will be well approximated by a Gaussian distribution with the same mean and variance and its standard deviation-to-mean ratio will be

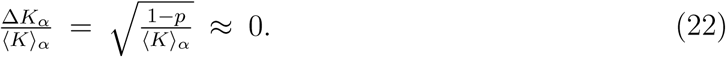

This makes it reasonable to approximate, in the terms of order 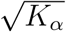 that appear in the expressions for *μ* and *σ, K*_*α*_ by its average ⟨*K*⟩_*α*_. This yelds

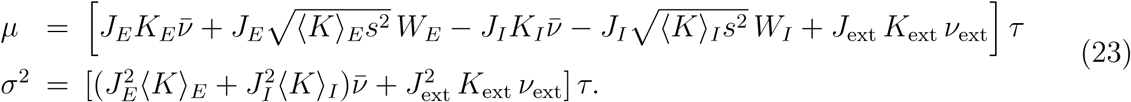

Under this approximation, *σ* can be regarded as a network constant and the expected value of *μ* is given simply by Eq. (9). Also, given that both the degrees *K*_*α*_ and the *W* s are independent Gaussian random variables, the quenched variability in the mean input *μ* across the network can be expressed by a single Gaussian random variable with combined variance

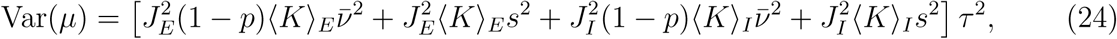

which is identical to Δ^2^ given by Eq. (10). Thus, we recover the formulation for the Erdös-Rényi case defined by Eqs. (8), (9), (10).

### B. Networks with correlated in- and out-degrees

In the previous section we have pointed out the dependency of the firing rate of a given neuron on its “characteristic” variables *K*_*αE*_, *K*_*αI*_, *W*_*αE*_, *W*_*αI*_. The result is general for any connectivity structure in which typical degrees are large but small compared with the system’s size *N*. Now we want to introduce the effect of having a non-zero correlation between individual in- and out-degrees.

To simplify the arguments, let us consider again the case of a single neuronal population. The variable *K* is the in-degree of the neuron under study, whereas *W* represents the variability in the sum of the firing rates of its pre-synaptic neurons. The theory presented here supposes that this sum originates from taking many independent realizations of the distribution of firing rates in the network. Since the number of elements that contribute to the sum is large, it is sufficient to know what the mean and variance of this distribution are. There does not seem to be a role for the out-degree in this formulation.

However, the distribution of firing rates within the set of pre-synaptic neurons to a given neuron can deviate from the distribution of firing rates in the network if in- and out-degrees are correlated, as we will explain now. Let us consider the process of taking one neuron at random and then picking one of its in-neighbors at random. Now we focus on the out-degree of the last neuron. This out-degree, as a random variable, does not follow the same distribution as the out-degrees in the network. For example, the network could potentially have many neurons with zero out-degree, but pre-synaptic neurons have out-degree of at least 1. Therefore, the distribution of out-degrees of the in-neighbors of a given neuron is biased. Now we consider the same random process but we look at in-degrees. The result is clear: if in/out-degrees are independent, the distribution will match the in-degree distribution in the network. On the contrary, if there is a correlation between degrees, the previous bias will be inherited by the in-degree distribution. Hence, the pre-synaptic neurons to a given neuron have in-degrees which follow a distribution that does not necessarily match the network distribution. Since the firing rate of a neuron is a function of its in-degree, this translates into a bias in terms of the *firing rates* of the pre-synaptic neighbors.

This bias can be analytically determined. Treating the degrees as discrete variables, the probability that the firing rate of a pre-synaptic neuron *j* to neuron *i* lies within the range (*v, v* + *δ*) once we know the in-degree of *i* is

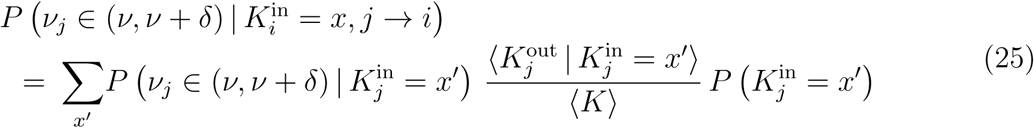

(see the Appendix for details). This result indicates that the distribution of firing rates among the pre-synaptic neurons of a given neuron does not necessarily follow the distribution of firing rates in the network. The bias is due to the fact that the expectation of the out-degree of a neuron *conditioned* on its in-degree is not necessarily equal to the expected out-degree. The ratio between this conditional expectation and the expectation itself is what alters the distribution of firing rates. Of course, when the degrees are independent this ratio is 1 and we recover the distribution of firing rates in the network. Notice, also, that the biased distribution is independent of the in-degree of the post-synaptic neuron, 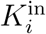, which means that the distribution of firing rates within the pre-synaptic neighbors of a given neuron is a *network* property (rather than a property associated with each post-synaptic neuron).

As a consequence, the sum of the rates of the pre-synaptic neurons to a given neuron (the variable *S* defined in (15)) is, in fact, a sum over the *biased* distribution of firing rates. We denote this new distribution with a star (*), so that 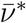 and *s*^*\**^ refer to the mean and the standard deviation of the biased version of the firing rate distribution.

The mean-field equations are the same as before with the difference that we have to replace 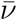 and *s*^2^ by 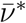 and (*s*^*\**^)^2^ in the definition of *μ* and *σ*^2^ (Eq. (16)). Analogously, in Eq. (20) 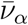 and 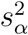 are replaced by 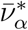 and 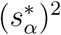, respectively. To self-consistently close the equations, we take into account the definition of the biased rate moments:

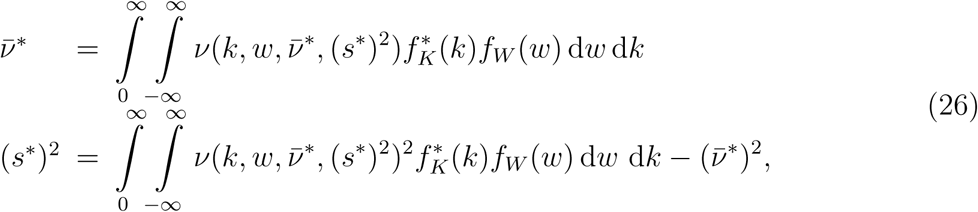

where 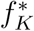 is the probability density function of the *biased* in-degree:

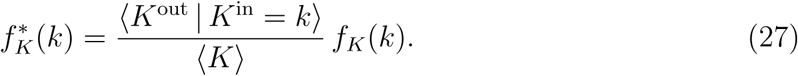

Interpreting the degrees as continuous variables, the conditional expectation is computed as

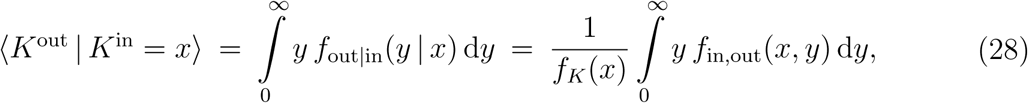

where *f*_out*|*in_ is the probability density of the out-degree conditioned to the in-degree and *f*_in,out_ is the probability density of the joint degree distribution. Analogous equations for the moments of the rate distributions can be obtained in the two-population scenario described by (20).

In all cases, the full firing rate distribution can be expressed with an integral of the form of Eq. (19). An alternative method for solving for the firing rate distribution is the following. *n* pairs of random variables (*k, w*) are generated from the distributions *f*_*K*_(*k*) and *f*_*W*_ (*w*). For each pair the steady-state firing rate is just *v* = *ϕ* (*μ*(*k, w*)*, σ*(*k*)). For large enough *n*, the histogram of firing rates generated in this way will be a good aproximation to the true distribution given by Eq. (19) or an analogous integral formulation. This is the method we use to generate the theoretical curves in Figs. 2 and 4.

We provide some examples in the next section.

### C. Examples

#### E/I network with ER connectivity for E→I, I→E and I→I connections and arbitrary degree distribution within the EE subnetwork

We first analyze the case of a network composed of excitatory (E) and inhibitory (I) neurons where connections from/to I neurons are generated independently with probability *p* (i.e., they have ER-like structure). The connectivity within the EE subnetwork, on the contrary, is created according to a joint in/out-degree distribution with correlation *ρ*. We assume that the number of external inputs that each neuron receives, *K*_ext_, is constant across the entire population.

We denote by ⟨*K*⟩_*αβ*_ and Δ*K*_*αβ*_ the mean and the standard deviation of the in-degree of a neuron in population *α* from population *β*. Since the ER in-degrees have a mean much larger than its standard deviation, the contributions of Δ*K*_*αβ*_ in expressions of the type 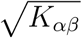 in Eq. (20) can be neglected for (*α, β*) ∈ {(*E, I*), (*I, E*), (*I, I*)} (and thus 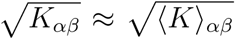 in these cases). On the other hand, and representing again the degrees by continuous variables, the remaining random variables *K*_*αI*_, *W*_*αE*_ and *W*_*αI*_ of Eq. (20) are independent and follow approximately Normal distributions, so they can be grouped together under a single Gaussian random variable *W*_*α*_ (this holds from the fact that the ER degrees follow Binomial distributions, which can be approximated by Gaussians in the large *N* limit). In the I population, the *K*_*IE*_ variable is also independent of the others and Gaussian, so it can be grouped with the other three variables through a new Gaussian variable *W*_*I*_.

This finally implies that the firing rates of neurons in the E population are parametrized by two quenched random variables (the excitatory in-degree *K*_*EE*_ and *W*_*E*_ ∼ *N* (0, 1)) and the firing rates in the I population, by a single variable (*W*_*I*_ ∼ *N* (0, 1)) which includes all the contributions coming from the degree and the rate heterogeneity.

We thus obtain

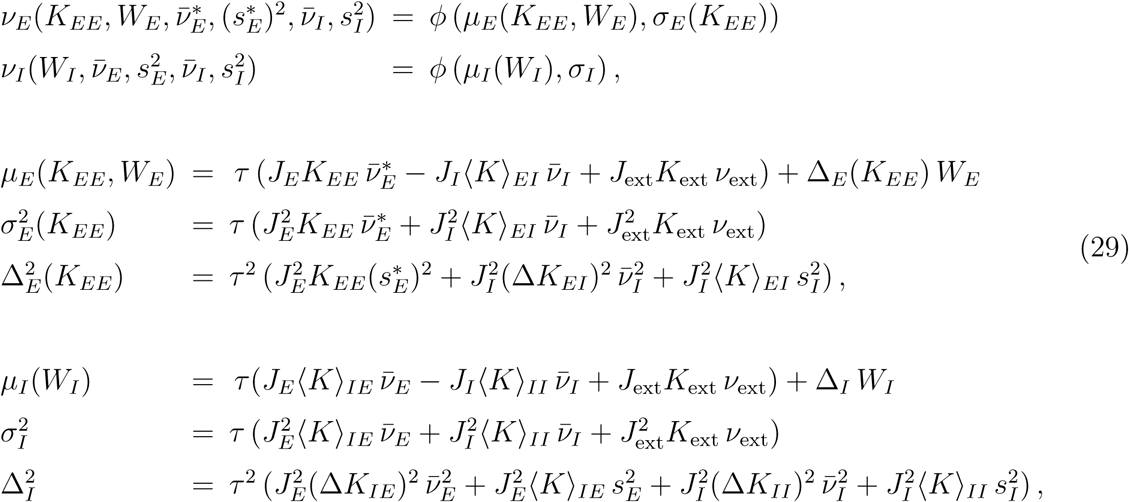

where *W*_*E*_*, W*_*I*_ ∼ *N* (0, 1).

Now the unknowns are 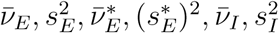. The equations that close the system are their definitions:

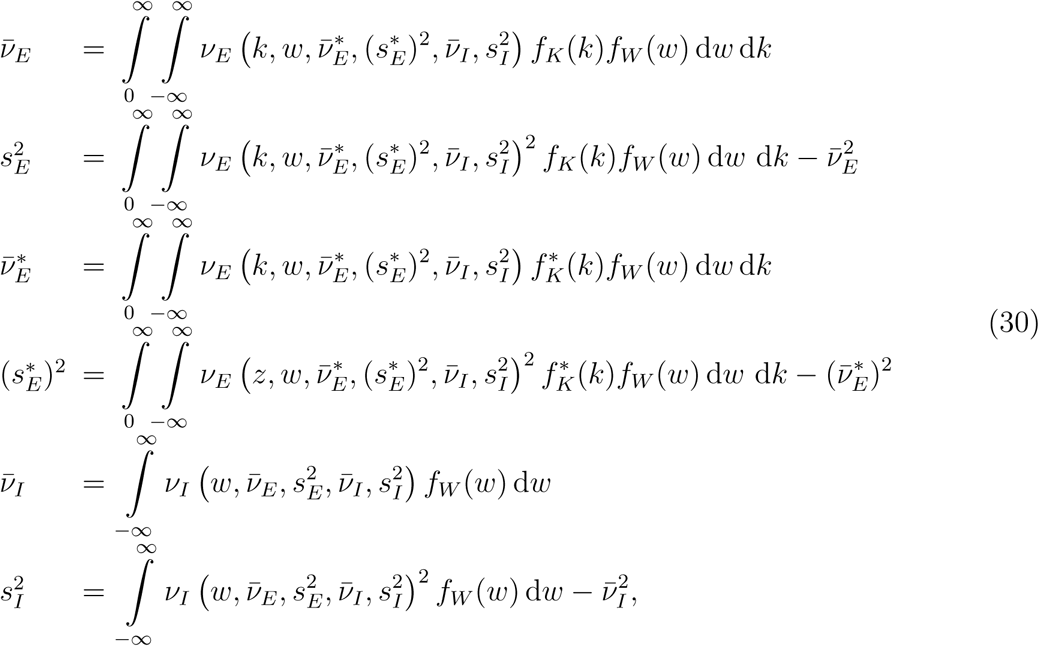

where *f*_*W*_, *f*_*K*_ and 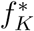 are the probability densities of a standard Gaussian random variable (*f*_*W*_), of the excitatory in-degree of E neurons (*f*_*K*_) and of the excitatory in-degree of E neurons which are pre-synaptic to a given E neuron 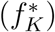. The last density depends on the joint in/out-degree distribution within the EE subnetwork *f*_in,out_ through

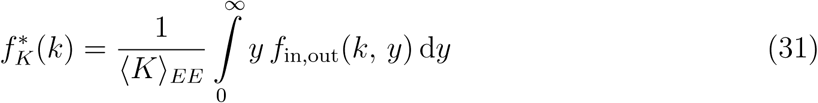

(this is just the result of putting together Eqs. (27) and (28)).

#### E/I network with an arbitrary degree distribution within the EE subnetwork and selective inhibition

In the preceding example, degrees within the EE subnetwork followed an arbitrary distribution, whereas the remaining connectivity was purely random (in the ER sense). But networks in which the EE in-degree distribution is much broader than in ER counterparts can present a clear unbalance in the inputs that E neurons receive [15, 18]. This feature is thought to be unrealistic because in physiological situations excitation and inhibition tend to cancel each other in the mean, leading to what has been called a *balanced state* [29–31].

The structure of connections involving inhibitory neurons has been less studied than that of excitatory-to-excitatory synapses. In very general terms, and despite a wide variety of interneuron types, inhibitory connectivity in the mammalian cortex appears to be denser, less specific and more homogeneous than connectivity between pyramidal neurons [32–34]. This makes ER models a reasonable preliminary framework to represent connections from/to inhibitory neurons. On the other hand, experiments on cortical slices have shown that the magnitude of the inhibitory component of the input adapts to that of the excitatory component through plasticity mechanisms that modulate the strength of inhibitory connections [29]. Such adaptations could be responsible for maintaining a proper balance between excitation and inhibition at the single cell level [15].

We consider here a situation in which the connectivity within the II subnetwork and from the E to the I population is of ER type but where the connections from I to E appear independently with a probability that depends (linearly) on the excitatory in-degree of the post-synaptic neuron, so as to compensate for the excess of excitation that some E neurons might receive. In this case, the inhibitory in-degree of an E neuron with excitatory in-degree *K*_*EE*_ follows, as before, a Binomial distribution whose parameters depend on *K*_*EE*_. This means that the same approximations that we made in the previous section are still valid now. The only new ingredient is the fact that ⟨*K*⟩_*EI*_ and Δ*K*_*EI*_ are not constant but functions of *K*_*EE*_:

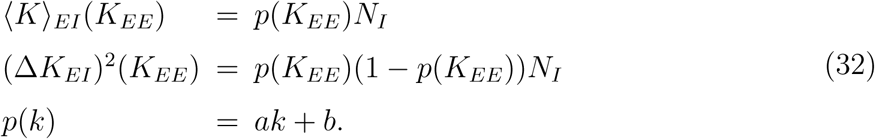

### D. Comparison between mean-field theory and computer simulations

We next compare the theory developed in the previous sections with computer simulations of networks of LIF neurons. We have chosen for illustration networks in which the EE degrees follow (integer versions of) Gaussian and Gamma distributions. We explore first the case of distributions with small variance and we move, afterwards, to networks with highly heterogeneous degree distributions.

#### Degree distributions with a low level of heterogeneity

We include in this category networks in which the degrees of the EE subnetwork follow distributions whose variance is close to the variance in ER networks of the same density. Since these networks have a low level of degree heterogeneity within the excitatory population, inhibition has been set to be completely homogeneous: connections from and to I neurons are created with a fixed probability *p* that is the same for the entire population.

We asked ourselves what the effect would be of varying the correlation between EE degrees and if the mean-field extension described earlier would be able to capture it. We started by imposing degrees in the EE subnetwork which are integer versions of Bivariate Normal distributions, keeping a relatively small marginal variance. Figure 2 shows three examples of networks that only differ in the correlation coefficient *ρ* between in- and out-excitatory degrees in the EE subnetwork (see Fig. 1A). The results indicate that the theory agrees well with simulations and is able to explain the modulation of the firing rate distribution as the parameter *ρ* is varied.

**FIG. 1.**
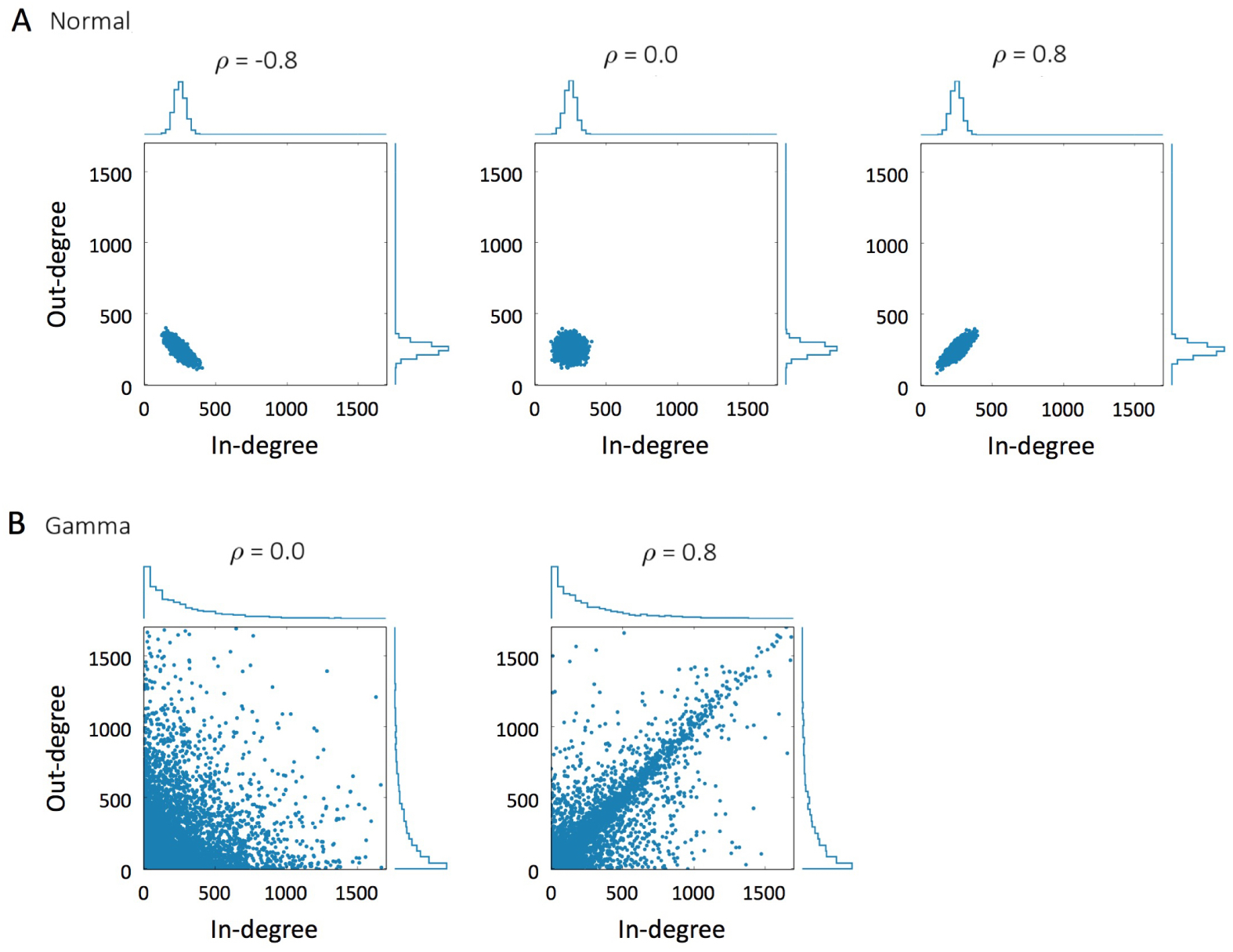
In/out-degree distributions within the excitatory subnetwork. **A** Networks whose degrees follow Normal(*µ, σ*) degree distributions with *µ* = 250, *σ* = 40 and correlation coefficient *ρ* = −0.8 (left), *ρ* = 0 (middle) and *ρ* = 0.8 (right). **B** Networks with Gamma(*κ, θ*) degree distributions, *κ* = 0.8, *θ* = 312.5 and correlation coefficient *ρ* = 0 (left) and *ρ* = 0.8 (right). In all the cases *N*_*E*_ = 5000.

**FIG. 2.**
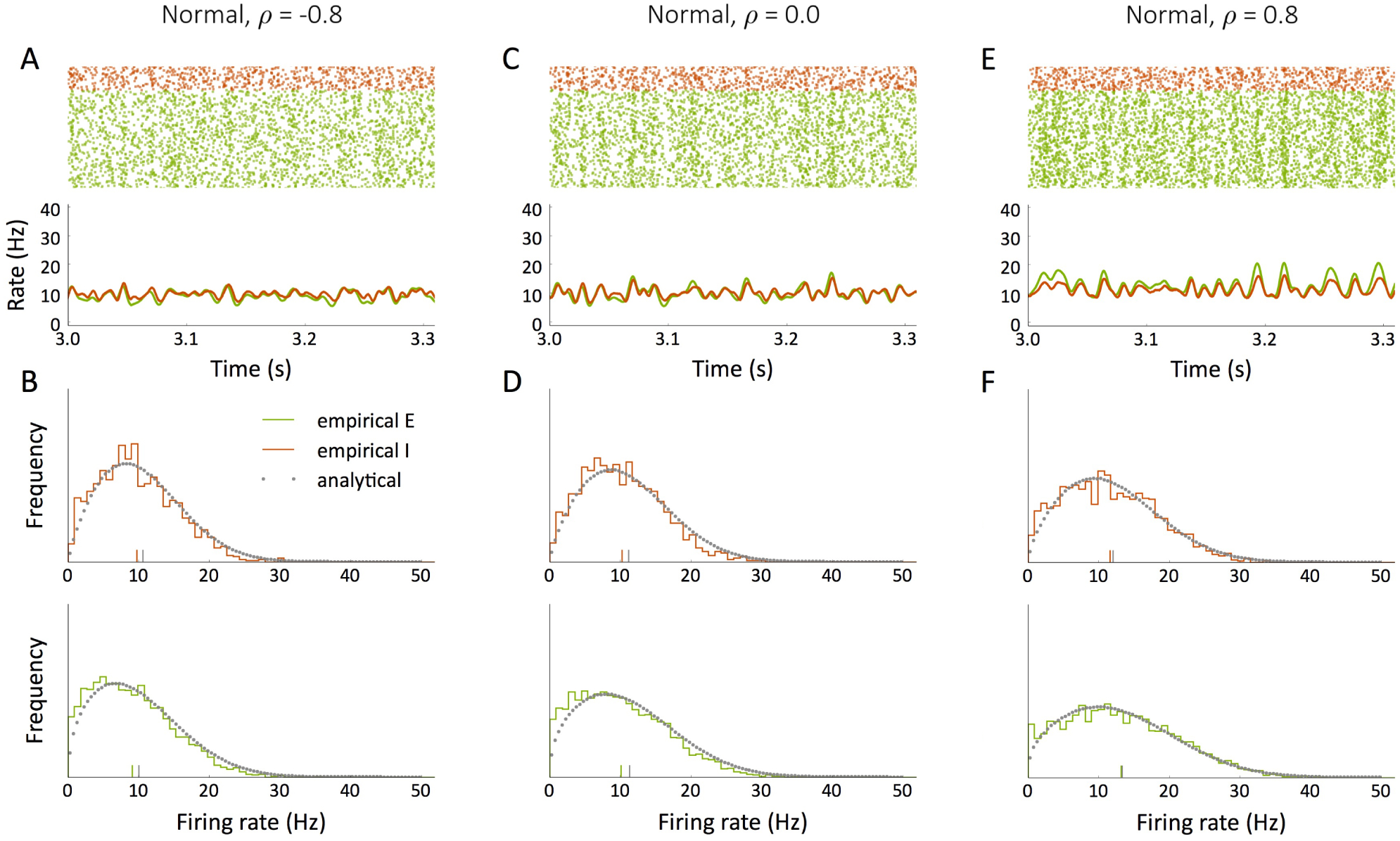
Dynamics of networks which only differ in the correlation coefficient *ρ* between in- and out-degrees within the EE subnetwork. In all the cases, the EE degree distribution is Normal(*µ, σ*) with *µ* = 250 and *σ* = 40, as in Fig. 1A. All the other connections are created independently with probability *p* = 0.05. **A** Raster plots and population firing rates of inhibitory (red) and excitatory (green) neurons when *ρ* = −0.8. **B** Firing rate distribution in the stationary state for I (red) and E (green) neurons, from simulations (continuous histogram) and from the analytical formula (grey dots) when *ρ* = −0.8. **C** Same as A for *ρ* = 0. **D** Same as B for *ρ* = 0. **E** Same as A for *ρ* = 0.8. **F** Same as B for *ρ* = 0.8. Vertical lines indicate empirical and analytical averages. In all the cases *N*_*E*_ = 5000, *N*_*I*_ = 1250, *τ* = 20 ms, *V*_*r*_ = 10 mV, *V*_*θ*_ = 20 mV, *τ*_*r*_ = 2 ms, *J*_*E*_ = 0.11 mV, *J*_*I*_ = *gJ*_*E*_, *g* = 8, *K*_ext_ = 1000, *v*_ext_ = 8.1 Hz, *J*_ext_ = 0.14 mV, *d*_*j*_ = 1.5 ms for all *j*.

#### Highly heterogeneous degree distributions

So far, we have shown that the presented theory gives an accurate prediction of the distribution of firing rates when the structure within the EE subnetwork is moderately heterogeneous. We next studied if the same equations can explain the behavior of the system when the structure is perturbed even more. We consider extreme cases of heterogeneous networks by using EE degrees which follow (integer versions of) Gamma(*κ, θ*) distributions with correlation coefficient *ρ*. Figure 1B shows two example EE degree distributions of this type. As noted before, in these cases we need to introduce a compensatory mechanism to keep a balance between excitatory and inhibitory inputs. We do so by creating I*→*E connections with a probability that depends on the excitatory in-degree *K*_*EE*_:

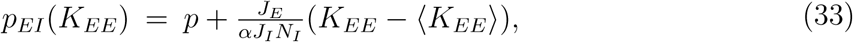

where *p* is the probability with which E*→*I and I*→*I connections are defined and *α* is a parameter which modifies the width of the distribution of I*→*E connection probabilities as shown in Figure 3.

**FIG. 3.**
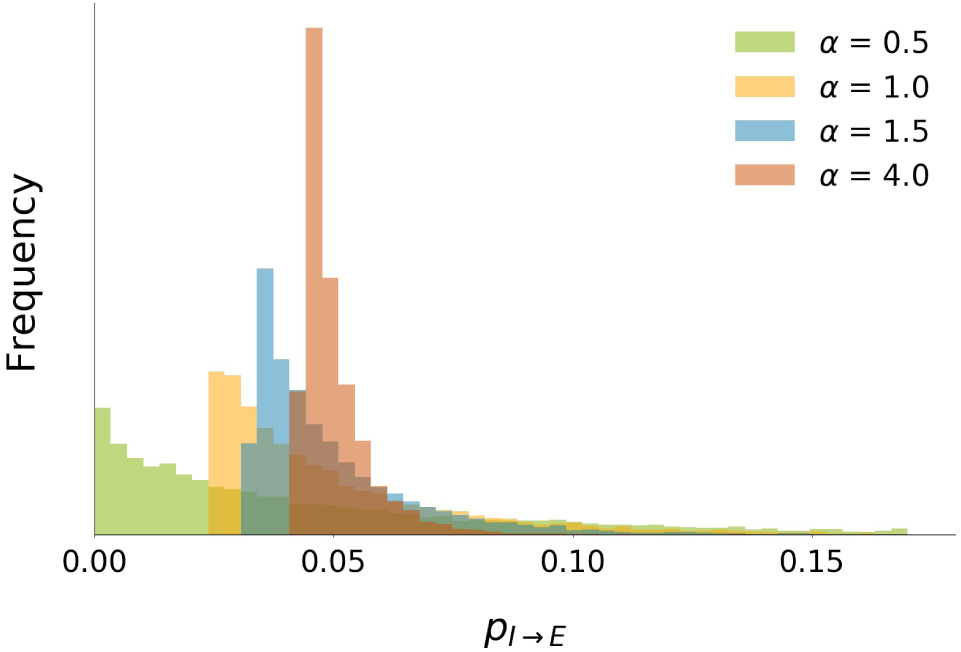
Distribution of I*→*E connection probabilities as a function of parameter *α* (see Eq. (33)) when the degree distribution within the EE subnetwork is highly heterogeneous (Gamma(*κ, θ*) with *κ* = 0.8, *θ* = 312.5 and *N*_*E*_ = 5000, as in Fig. 1B).

Again, the theory predicts the observed firing rate distribution with good accuracy. In the examples of Fig. 4 (whose structures correspond to those in Fig. 1B), the variation of the correlation coefficient *ρ* from 0 to 0.8 has a dramatic effect on the firing rates in the network, especially in the E population (Fig. 4B,D). Notice that the two networks of this example only differ in *ρ*; their marginal degree distributions are statistically identical. This shows the importance that the degree correlation has on dynamics. The presence of neurons which *at the same time* receive many inputs from the network and have an influence on many other neurons largely influences the dynamical properties of the network as a whole. Moreover, this effect of *ρ* is almost perfectly captured by the introduction of the moments of the biased rate distribution, 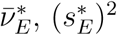 in the analytical formalism.

**FIG. 4.**
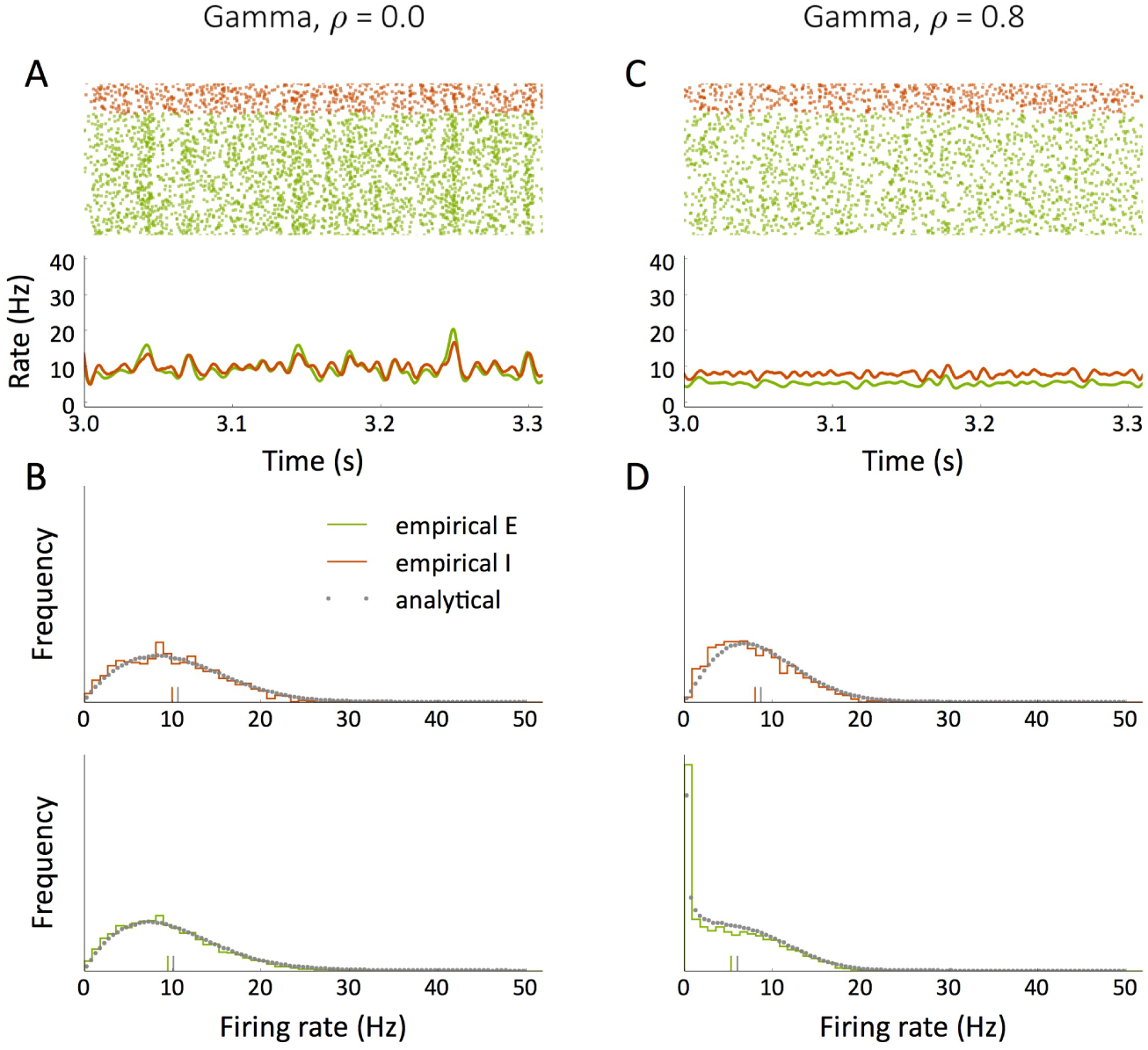
Dynamics of networks which only differ in the correlation coefficient *ρ* between in- and out-degrees within the EE subnetwork. In all the cases, the EE degree distribution is Gamma(*κ, θ*) with *κ* = 0.8 and *θ* = 312.5, as in Fig. 1B. **A** Raster plots and population firing rates of inhibitory (red) and excitatory (green) neurons when *ρ* = 0. **B** Firing rate distribution in the stationary state for I (red) and E (green) neurons, from simulations (continuous histogram) and from the analytical formula (grey dots) when *ρ* = 0. **C** Same as A for *ρ* = 0.8. **D** Same as B for *ρ* = 0.8. Vertical lines indicate empirical and analytical averages. In all the cases *N*_*E*_ = 5000, *N*_*I*_ = 1250, *p* = 0.05, *τ* = 20 ms, *V*_*r*_ = 10 mV, *V*_*θ*_ = 20 mV, *τ*_*r*_ = 2 ms, *J*_*E*_ = 0.11 mV, *J*_*I*_ = *gJ*_*E*_, *g* = 8, *α* = 1, *K*_ext_ = 1000, *v*_ext_ = 8.1 Hz, *J*_ext_ = 0.14 mV, *d*_*j*_ = 1.5 ms for all *j*.

## IV. A POSSIBLE FUNCTIONAL ROLE OF DEGREE CORRELATIONS

In the previous sections we studied the role of *ρ* in shaping the repertoire of firing rates in a stationary, asynchronous state. We next asked if the presence of “in/out-hubs” in networks with positive *ρ* and large degree heterogeneity could modify the response to transient stimuli compared with networks with independent in- and out-degrees.

To do so, we applied transient pulses of stimulation to a fraction *f*_ext_ of excitatory neurons, chosen at random. The modified version of the voltage dynamics for a stimulated neuron *i* is, for *t* within the stimulation period,

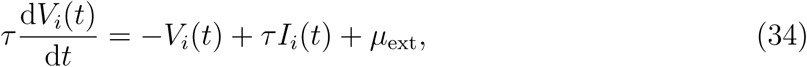

where *I*_*i*_(*t*) takes into account all the inputs coming from other neurons (including the external ones) and *µ*_ext_ is the magnitude of the transient stimulation.

When the fraction of stimulated neurons is small, there is no response apart from a tiny increase in population rates while the stimulus is applied, independently of the network’s topology. This behavior persists regardless of *f*_ext_ as long as the EE degrees are not correlated. However, for large enough positive values of *ρ*, the response changes dramatically. In this case there is a critical fraction of stimulated neurons above which the network transiently enters in a qualitatively distinct dynamical regime, characterized by bursting and synchronous activity. Such a “responsive” state lasts even after the stimulus has been removed, although the network eventually relaxes back to its original asynchronous state, as shown in Fig. 5.

**FIG. 5.**
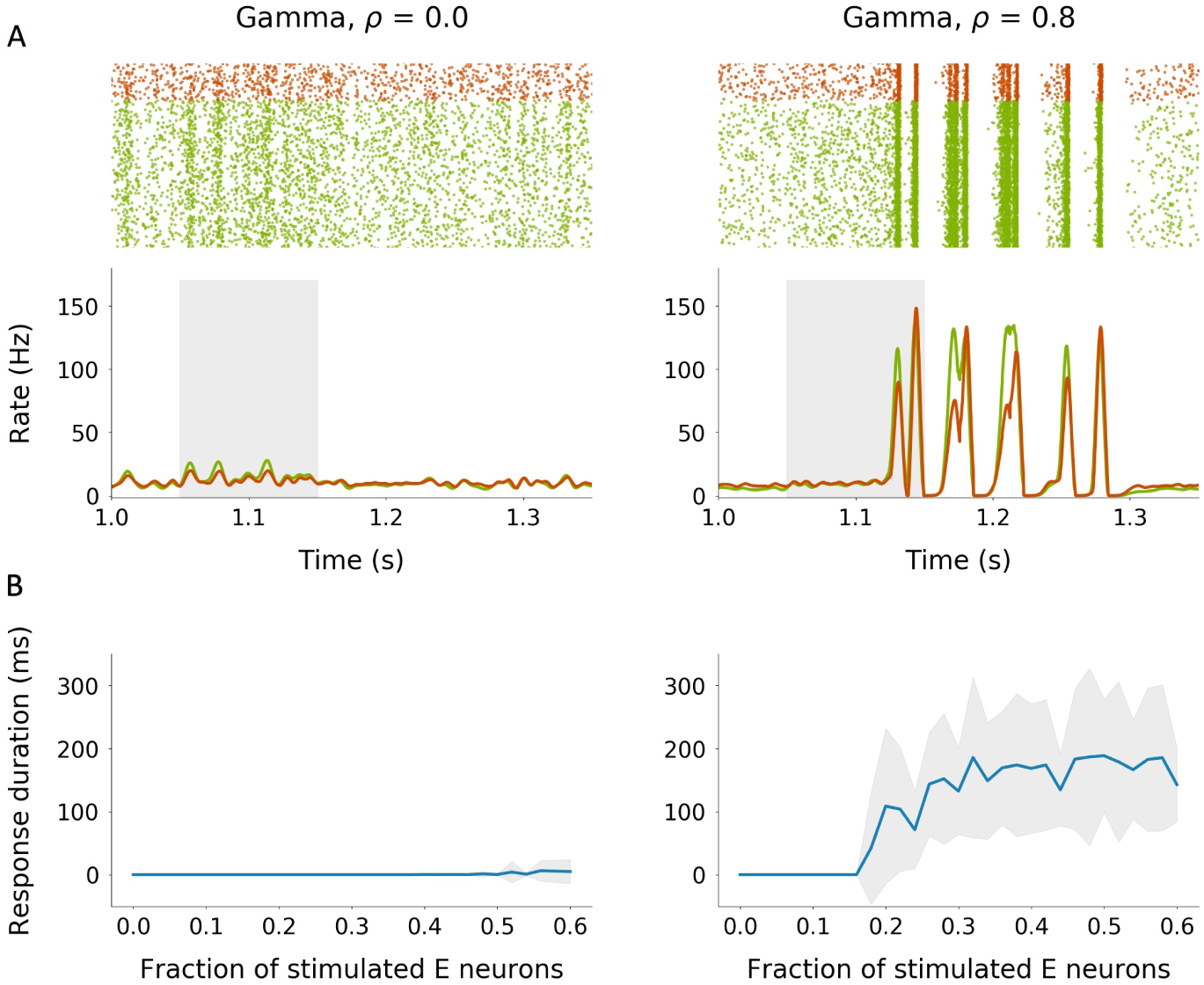
Response to transient stimulation of two different kinds of topologies for the EE subnetwork: broad Gamma distribution without correlation (left), and broad Gamma distribution with positive correlation (right). **A** Raster plots and population firing rates for a stimulation of mag-nitude *µ*_ext_ = 5 mV applied to a fraction of *f*_ext_ = 0.2 E neurons during *t*_ext_ = 100 ms (shaded region). **B** Total duration of the network’s response (average *±* standard deviation) as a function of *f*_ext_ for *µ*_*ext*_ = 5 mV and *t*_ext_ = 100 ms. The mean and the standard deviation for each *f*_ext_ have been computed over 20 simulations in which the stimulated neurons are the same. The remaining parameters are the same as in Fig. 4.

The region of burst-like responses occurs in the asynchronous regime of the network activity, but near an instability to spontaneous population bursting. This is shown in the phase diagram, Fig. 6, in which we varied the correlation between in- and out-degrees in the EE subnetwork (*ρ*) and the factor *α*, which controls the width of the distribution of I*→*E connection probabilities, see Fig. 3. Bursting occurs spontaneously in the blue region of Fig. 6. Near this boundary, but in the asynchronous regime, bursting can be induced by transient stimuli (see, for example, the configuration indicated by the green star, corresponding to the network of Fig. 5B). This occurs mainly for large values of *ρ*, which again suggests that positive degree correlations facilitate the responsiveness of the network to transient stimuli, and seems to be a prerequisite for these networks to detect transient external signals.

**FIG. 6.**
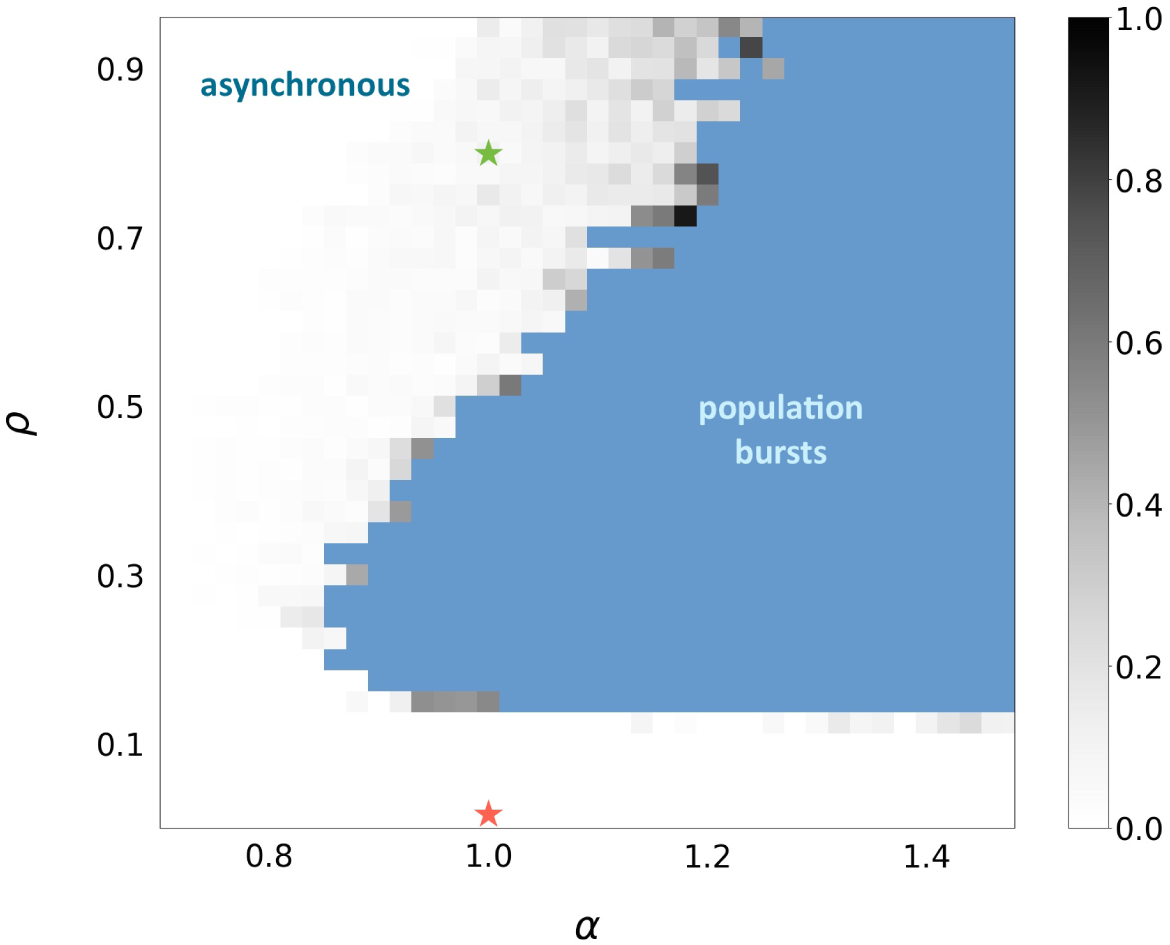
Ability of the network to exhibit bursting activity when the EE subnetwork has a broad Gamma degree distribution as a function of the correlation coefficient between EE degrees (*ρ*) and the narrowness of the I*→*E connection probability distribution (*α*). The blue region corresponds to configurations which exhibit spontaneous population bursts in the absence of transient stimulation. The other configurations are not able to exhibit these bursts unless they receive a transient stimulus. In these cases, the duration of the bursts once the stimulus has been removed is shown in a gray color scale, relative to the maximum oberved burst, which was 1848ms. Red and green stars indicate the configurations corresponding to the networks of Figure 5A and B, respectively. Each square corresponds to an average of 10 independent simulations on the same network realization. The criterion to consider a configuration in the “population bursts” group is that the average relative duration of the spontaneous bursts be larger than 0.15. The magnitude of the transient stimulus was *µ*_ext_ = 5 mV and it was given to a fraction *f*_ext_ = 0.3 of E neurons during *t*_ext_ = 100 ms. The remaining parameters are the same as in Figs. 4 and 5.

Transient bursts, however, are not found in networks with a low level of degree hetero-geneity. For example, networks with Normal degree distributions and degree variance as in Fig. 1A do not exhibit this behavior, as shown in Fig. 7.

**FIG. 7.**
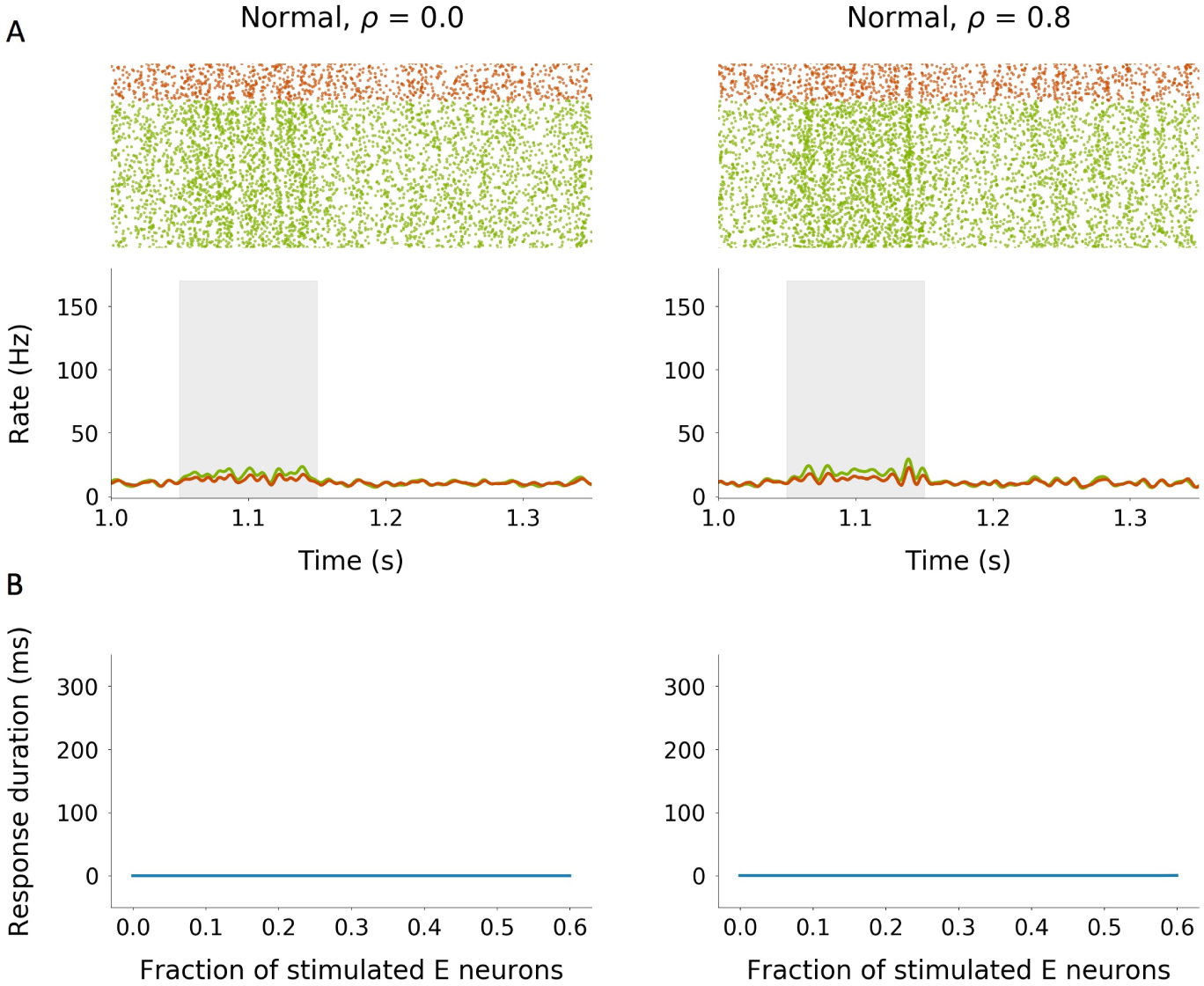
Response to transient stimulation of two different kinds of topologies for the EE sub-network: narrow Normal distribution without correlation (left), and narrow Normal distribution with positive correlation (right). **A** Raster plots and population firing rates for a stimulation of magnitude *µ*_ext_ = 5 mV applied to a fraction of *f*_ext_ = 0.2 E neurons during *t*_ext_ = 100 ms (shaded region). **B** Total duration of the network’s response (average *±* standard deviation) as a function of *f*_ext_ for *µ*_*ext*_ = 5 mV and *t*_ext_ = 100 ms. The mean and the standard deviation for each *f*_ext_ have been computed over 20 simulations in which the stimulated neurons are the same. The remaining parameters are the same as in Fig. 4.

Therefore, the degree-correlation *ρ* is important in defining how external inputs are prop-agated through the network. The presence of a significant number of in/out-hubs likely enhances the transmission of information because such nodes act as “organizing centers”: they receive information from a large fraction of the network and, simultaneously, transmit it to a large number of other units.

## V. DISCUSSION

In this paper we have extended a well-known mean-field formalism [24] to a more general family of possible connectivities. The theory presented here does not exclude the common topologies, but can be applied to any network whose structure is determined solely by its degree distribution. Therefore, homogeneous and purely random (ER) models are still included in the formalism. It is important to bear in mind, however, that the theory does not apply to networks whose structure is determined by other principles: despite the fact that any network family has a characteristic degree distribution, this does not necessarily imply that the distribution *per se* defines its structure.

Briefly, neurons in heterogeneous networks can be parametrized by a (random) variable which defines their “rate identity” in the network. The rate of every neuron is then a function of this variable, which captures the neuron-to-neuron differences in terms of both the number of inputs received and the rates of the input units. In classical ER models, such a variable is a scalar and can be assumed to be normally distributed (because the in-degrees follow approximately Gaussian distributions). The new ingredient when dealing with a network whose in-degrees follow an arbitrary distribution is that the above mentioned neuron-to-neuron variability is no longer captured by a scalar but by a two-component vector. This is so because the two contributions to the neuronal variability-differences in in-degree and differences in the firing rates of input neurons- can no longer be grouped into a single variable. Furthermore, when in-degrees and out-degrees are correlated, the set of all pre-synaptic inputs to a neuron represent a biased firing rate distribution compared to the full, network-wide distribution. The reason is simply that a neuron is more likely to receive an input from another neuron with high out-degree. When degree-correlations are present, this bias is carried over to the in-degree, and hence to the firing rate. The effect of such a rate bias can be analytically computed and introduced in the mean-field equations. The final mean-field formulation takes into account not only the moments of the firing rate distribution but also those of the biased rate. The macroscopic unknowns of the system are, then, the mean and variance of the rate distribution and also the moments of the biased rate distribution, and the equations can be self-consistently closed by using the definitions of these four quantities.

There is experimental evidence that firing rates *in vivo* follow skewed, log-normal distributions [2, 3]. This has been proposed to be the result of a Normal distribution of inputs (which occurs, for example, when in-degrees follow approximately Normal distributions) plus a non-linearity of the input-rate function [26]. However, recent studies suggest that the in-degrees of different neurons could be more heterogeneous than previously expected [14, 15]. The mean-field extension presented here can help to quantitatively explore the relationship between the degree heterogeneity and the resulting distribution of firing rates. It could even be used to address the inverse problem, that is, deriving properties of the underlying topology given some (experimentally observed) constraints in the repertoire of firing rates.

The role of the above-mentioned biased firing rate goes beyond its technical use in the described formalism. Our observations show that, in fact, the relevant parameter when studying the behavior of the system is precisely this biased distribution. What neurons really “perceive” is the firing rates in their set of neighbors, not in the entire network. A network might contain a large fraction of very active neurons which nevertheless do not project to any others. In terms of function, such neurons presumably do not play any role because they have no influence. Thus, the firing rate distribution restricted to the set of possible pre-synaptic neurons is a much more relevant dynamical magnitude. This suggests that determining the firing rate distribution in real networks may be not very informative unless it is accompanied by a thorough study of connectivity.

Finally, we explored the role of the degree-correlation *ρ* in shaping the spontaneous activity. In E-I networks with positive correlations in the recurrent excitatory degrees alone, and with ER connectivity in the remaining connections, the spontaneous activity is highly skewed and unrealistic: many neurons are silent and some fire at exceedingly high rates. In order to offset this skewness, we make the inhibitory-to-excitatory connection probability proportional to the recurrent excitatory in-degree. The inhibition is therefore “selective” to the excitatory in-degree. In such a network, the spontaneous activity is irregular and asynchronous even for very high, positive correlations, as long as the selective inhibition is sufficiently strong, see Fig. 6. When the selective inhibition is weaker, strong enough positive degree-correlations lead to a ongoing population bursts. For degree-correlations and inhibition selectivity which place the system in the asynchronous regime but near the boundary to bursting, an external stimulus can trigger a long-lasting, yet transient burst. This behavior resembles that of short-term memory networks, in which the memory is stored as an activity pattern that is maintained in time even in the absence of the stimulus that originated it [35]. Such observations could link degree correlations –which could be present in real cortical microcircuits [13]- with enhanced abilities to transmit and process external inputs.

## ACKNOWLEDGMENTS

M.V. has received financial support through the “la Caixa” Fellowship Grant for Post-Graduate Studies, “la Caixa” Banking Foundation, Barcelona, Spain. A.R. acknowledges grants BFU2012-33413 and MTM2015-71509-C2-1-R from the Spanish Ministry of Economics and Competitiveness and grant 2014 SGR 1265 4662 for the Emergent Group “Network Dynamics” from the Generalitat de Catalunya. This work was partially funded by the CERCA program of the Generalitat de Catalunya. The authors also thank Nicolas Brunel for useful discussions.

## APPENDIX: PRE-SYNAPTIC FIRING RATE DISTRIBUTION

First, let us notice that if *i* and *j* are two random neurons and 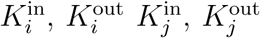 are their in- and out-degrees,

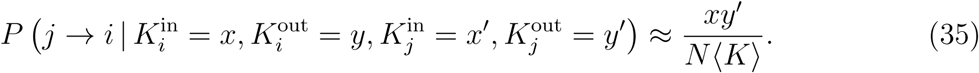

Now we compute the probability that the firing rate of a pre-synaptic neuron *j* to neuron *i* lies within the range (*v, v* + *δ*) once we know the in-degree of *i*:

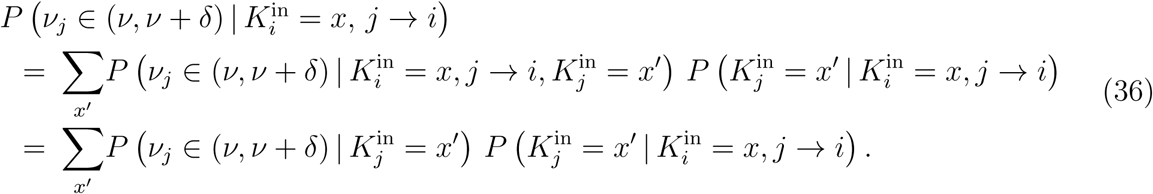

On the other hand,

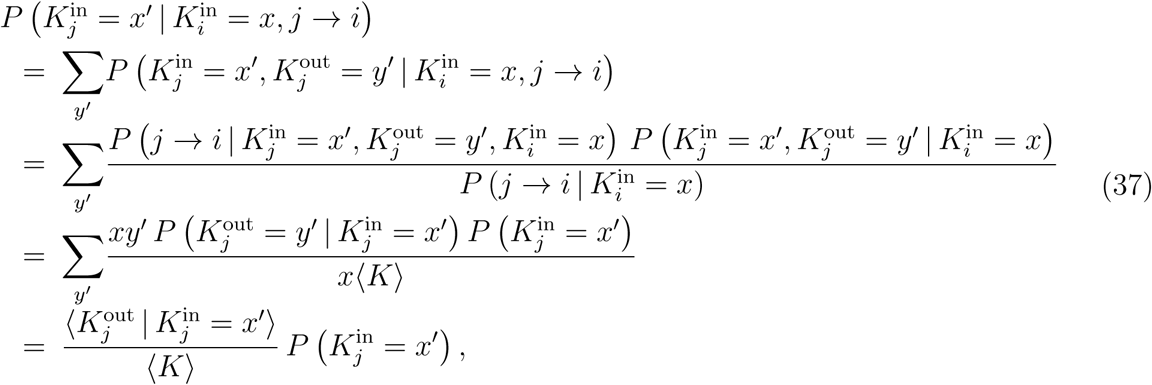

where we have used the fact that, in the considered networks, the degrees of different neurons are independent variables. Inserting (37) into (36) gives

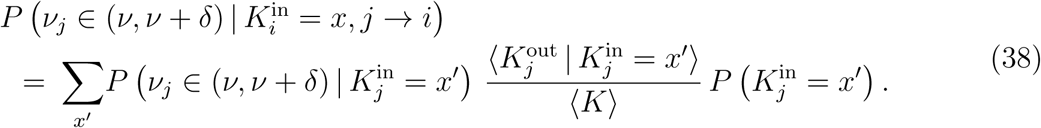

